# Ultrasound Mediated Delivery of Quantum Dots from a Capsule Endoscope to the Gastrointestinal Wall

**DOI:** 10.1101/2020.02.24.963066

**Authors:** Fraser Stewart, Gerard Cummins, Mihnea V. Turcanu, Benjamin F. Cox, Alan Prescott, Eddie Clutton, Ian P. Newton, Marc P.Y. Desmulliez, M. Thanou, H. Mulvana, Sandy Cochran, Inke Näthke

## Abstract

Biologic drugs, defined as therapeutic agents produced from or containing components of a living organism, are of growing importance to the pharmaceutical industry. Though oral delivery of medicine is convenient, biologics require invasive injections because of their poor bioavailability via oral routes. Delivery of biologics to the small intestine using electronic delivery with devices that are similar to capsule endoscopes is a promising means of overcoming this limitation and does not require reformulation of the therapeutic agent. The efficacy of such capsule devices for drug delivery could be further improved by increasing the permeability of the intestinal tract lining with an integrated ultrasound transducer to increase uptake. This paper describes a novel proof of concept capsule device capable of electronic application of focused ultrasound and delivery of therapeutic agents. Fluorescent markers, which were chosen as a model drug, were used to demonstrate in-vivo delivery in the porcine small intestine with this capsule. We show that the fluorescent markers can penetrate the mucus layer of the small intestine at low acoustic powers when combining microbubbles with focussed ultrasound. These findings suggest that the use of focused ultrasound together with microbubbles could play a role in the oral delivery of biologic therapeutics.

## Introduction

Oral delivery of therapeutic agents is generally the preferred route of administration due to increased patient acceptance^1^ and convenience compared to parenteral routes. Many pharmaceuticals, once swallowed, are absorbed in the gastrointestinal (GI) tract, usually in the small intestine, which has greater absorptive capacity than other parts of the GI tract due to factors such as its length and surface area of up to 6 m and 200 m^2^ respectively in adults^2^. However, the challenging environment of the GI tract limits the successful absorption, and the ability to establish sufficient systemic levels of therapeutics. The pH along the gastrointestinal (GI) tract varies widely, and the gut contains many enzymes that reduce the stability, bioavailability, and thus effective delivery of many biomacromolecules^3,4^. There are also physical barriers to uptake that must be overcome before any biomacromolecular drugs can pass through the walls of the GI tract and reach the desired site in the body. First, the drugs must pass through the mucus that coats the intestinal epithelium^5^. Then, they must breach the barrier provided by the epithelial layer, specifically tight junctions between adjacent epithelial cells. These and other constraints have limited oral drug delivery to small molecules^4^. A more effective GI drug delivery system should achieve efficient delivery of a broader class of therapeutic agents, including biologics, with increased bioavailability and minimized toxic side effects and a reduction in the quantity that needs to be administered. Such a system should also remove the need for significant reformulations of the drugs and, for gastrointestinal diseases, enable localized treatment of conditions such as inflammatory bowel disease (IBD)^6^.

Pharmaceutical technologies such as multilayered tablets, intestinal patches, microneedle patches, hydrogels, and exosomes provide a means of controlling drug delivery in the intestine^7^. However, these methods may require the reformulation of therapeutic agents to guarantee compatibility with the chosen technique and ensure the efficacy of the drug^8^ though these methods are still not useful to ensure effective oral delivery of biologicals.

Another strategy for more effective oral drug delivery has emerged from advances in electronic miniaturization, specifically the development of capsule endoscopy (CE). Endoscopic capsules can be swallowed, and they contain a camera and associated electronic subsystems that allow optical imaging of the GI mucosa^9^. Clinical use of such capsules, primarily for the detection of occult GI bleeding, has increased since their introduction in the early 1990s^9^. The ability of CE to transit the entire GI tract makes them particularly suitable for imaging diseases that affect remote sections of the small bowel.

Capsule endoscopy, mainly used for its diagnostic advantages, is also widely recognized as a platform with therapeutic potential for the electronic delivery of commonly ingested drugs^3^. Capsule endoscopes could transport and release any drug to a region of the GI tract within a specified time after ingestion or upon detection of a change in pH. The clinical potential^10^ of such systems is illustrated by positive results obtained from clinical trials with existing drug delivery capsules such as Intellicap^11^, Intellisite^12^, and Enterion^13^. However, the potential of capsule devices for therapeutic applications goes further, as they could also be modified to improve the bioavailability of drugs by actively increasing tissue layer permeability. This approach would enhance the passage of therapeutic agents across tissue barriers to allow more efficacious treatments and reduction in doses that need to be delivered^4^.

Increased and reversible permeabilization of the tissue layers lining the GI tract can be induced through the use of ultrasound (US)^14–16^. Ultrasound-mediated targeted delivery was first considered in the 1980s as a method for targeting and enhancing therapeutic agent delivery^17^, with Phase III clinical trials currently underway for one of the first clinical treatments using this technique^18^. Conventional ultrasound-mediated targeted drug delivery (UmTDD) systems typically consist of an extracorporeally situated US transducer coupled to the skin with water or gel and aimed toward a target site^19^. Though this approach is not constrained by transducer size or power budget, the presence of bone or gas in the path of the US beam can produce unintended hotspots and shadows, and the patient must remain still to maintain the focus of the beam on the desired target.

US contrast agents, such as microbubbles (MBs), can be used to amplify these biophysical effects, enabling ultrasound-induced reversible permeabilization of cell membranes at low acoustic powers. Microbubbles consist of an inert gas core stabilized by a lipid or polymer shell typically 0.8-10 μm in diameter^20^. Several papers have demonstrated the ability of MBs to amplify the biophysical effects of ultrasound, such as cavitation. The gas-filled, compressible core of MBs makes them responsive to ultrasound, causing them to compress and expand alternately. This cyclical behavior can increase cell permeability due to the formation of pores caused by either the interaction between microbubbles and cell membranes at low acoustic pressures, referred to as stable cavitation, or through shockwaves generated by the collapse of microbubbles proximal to the cell membrane under high acoustic pressures, referred to as inertial cavitation^14^.

Some of the challenges associated with conventional UmTDD could be solved by placing the US transducer intracorporeally. This approach achieved a tenfold increase in the permeation of the anti-inflammatory drug 5-aminosalicylic acid when administered rectally^21^. However, in this case, the size of the device limited positioning to the rectum only.

Therapeutic intervention using UmTDD along the entire length of the GI tract could be achieved by placing the US source in a device resembling a CE. Such a device would also remove limitations caused by patient movement, obstruction by bone or gas associated with extracorporeal US. Intracorporeal UmTDD requires the miniaturization of the US transducer, resulting in lower power consumption, a reduction of the US intensity, and a decrease in the therapeutic efficiency of the treatment. However, the extent of these effects can be mitigated using MBs. The ability to increase cell permeability at low acoustic pressures due to the interaction between MBs and the cells is essential for the successful operation of an UmTDD capsule.

Following previous work^22–24^, this paper describes the design, manufacture, and characterization of a proof-of-concept therapeutic capsule for ultrasound-mediated delivery of agents using MBs. Previous work demonstrated that MBs, in conjunction with focused US reduced barrier function more effectively than insonation or MBs alone in Caco-2 cell monolayers^24^. Importantly, the decrease in TEER was temporary, and TEER to normal values was rapidly restored after insonation stopped. New results present the design of a new and more effective ultrasound-mediated delivery capsule that can successfully deliver fluorescent particles using MBs and insonation to *ex vivo* and *in vivo* tissues. The results of the *in vivo* experiments demonstrate that fluorescent markers can penetrate the mucus layer lining the small intestine, illustrating the potential for a new method of GI drug delivery.

## Results

### Ultrasound Transducer Physical Characteristics

A spherically focused US transducer with an outer diameter of 5 mm, a radius of curvature of 15 mm, and a central hole 1 mm in diameter was created using PZ26 piezoceramic material (Figure1a). The US was focused to create bioeffects in the focal region of the transducer. The central hole facilitated the integration of the MB/drug delivery channel and simplified this new capsule design^23^

**Figure 1:**
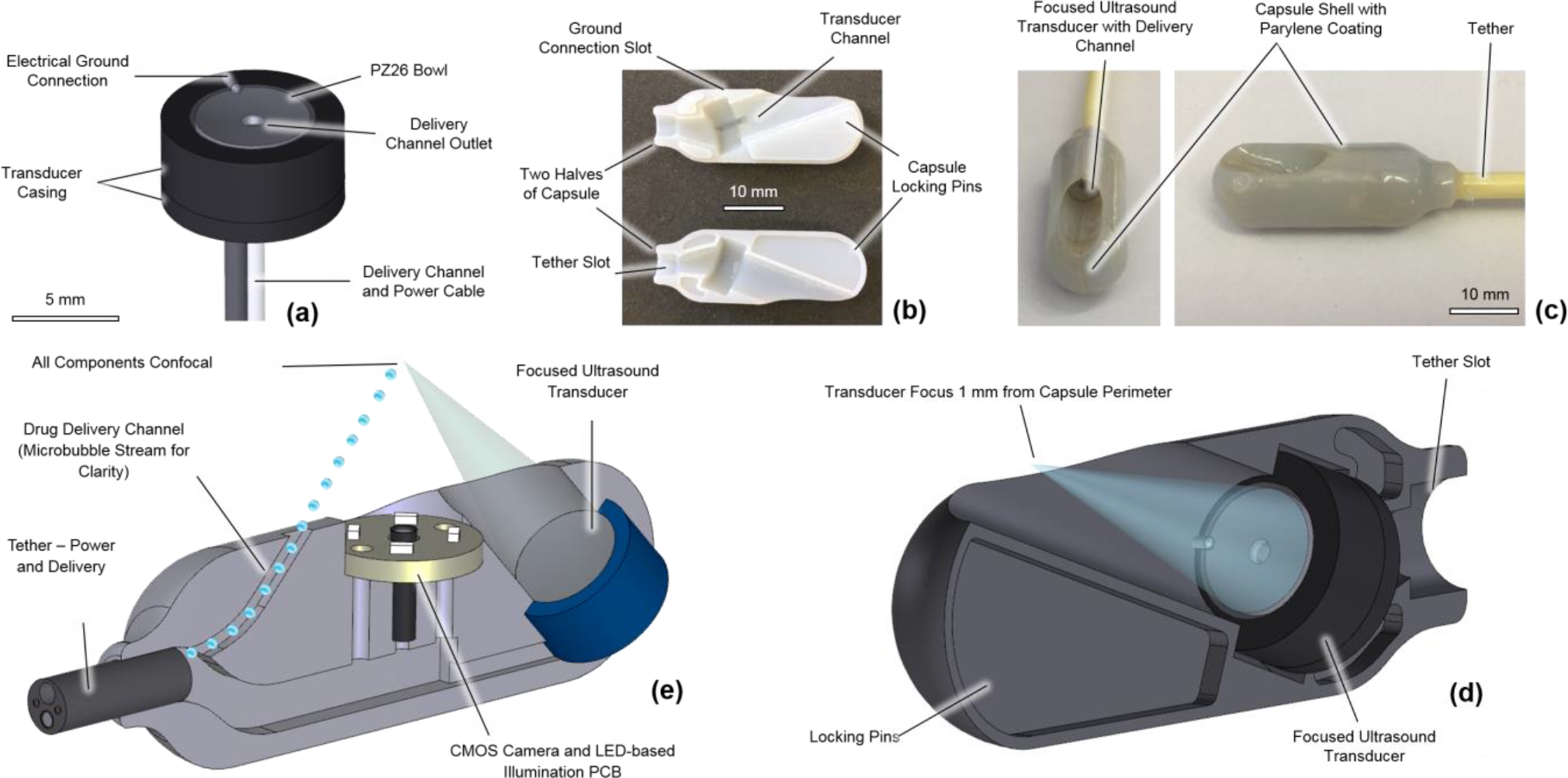
(a) Schematic of ultrasound transducer used in delivery capsule. (b) Locking mechanism for aligning capsule. (c) External view of capsule for *in vivo* drug delivery. (d) Cross-sectional image of *in vivo* delivery capsule. (e) Cross-sectional image of original prototype capsule^24^.

### Ultrasound Transducer Fabrication and Characterisation

Electrical impedance spectroscopy with a 4395A Impedance Analyzer (Keysight Technologies, Santa Clara, CA, USA) established that the central operating frequency of the transducers, when submerged in water, was 3.98 MHz. The magnitude of the electrical impedance at this frequency was within 10% of the electrical impedance of the attached power cable; therefore, no electrical matching was required. Acoustic output power measurements were performed with an input power (W_IN_) in the range of 20.1 mW to 223.6 mW. Results are shown in Table 1 and Table 2 for the transducers used for the *ex vivo* tissue and *in vivo* porcine experiments respectively, with a minimal difference in output power (W_OUT_) for the two transducers. The beam diameter of the transducer at −6 dB was measured using a commercial US field mapping (USFM) measurement system (Precision Acoustics Ltd., Dorchester, England).

**Table 1:** Output parameters of the miniature focused ultrasound transducer used for the *ex-vivo* tissue experiments

| **Voltage**<br>**(V<sub>pp</sub>)** | **W<sub>IN</sub>**<br>**(mW)** | **W<sub>OUT</sub>**<br>**(mW)** | **Efficiency**<br>**(%)** | **Acoustic Pressure**<br>**(kPa)** | **IAC**<br>**(Wcm<sup>-2</sup>)** |
| --- | --- | --- | --- | --- | --- |
| 3 | 20.1 | 9.6 ± 0.7 | 42.1 ± 3.3 | 344.5 ± 065.5 | 1.4 ± 0.0 |
| 4 | 35.8 | 20.2 ± 3.2 | 49.8 ± 3.9 | 443.4 ± 084.2 | 1.6 ± 0.1 |
| 5 | 55.9 | 31.3 ± 4.2 | 49.4 ± 3.9 | 544.7 ± 103.5 | 2.3 ± 0.1 |
| 6 | 80.5 | 42.4 ± 3.0 | 46.4 ± 3.7 | 636.3 ± 120.9 | 3.0 ± 0.2 |
| 7 | 109.6 | 56.8 ± 4.0 | 45.8 ± 3.6 | 741.3 ± 140.8 | 3.4 ± 0.2 |
| 8 | 143.1 | 76.6 ± 5.4 | 47.3 ± 3.7 | 826.8 ± 157.1 | 4.3 ± 0.3 |
| 9 | 181.1 | 92.5 ± 6.5 | 44.9 ± 3.6 | 917.4 ± 174.3 | 4.8 ± 0.4 |
| 10 | 223.6 | 115.1 ± 8.1 | 45.3 ± 3.6 | 1016.2 ± 193.1 | 6.1 ± 0.5 |

**Table 2:** Output parameters of the miniature focused ultrasound transducer used for the *in-vivo* capsule.

| **Voltage**<br>**(V<sub>pp</sub>)** | **W<sub>IN</sub>**<br>**(mW)** | **W<sub>OUT</sub>**<br>**(mW)** | **Efficiency**<br>**(%)** | **Acoustic Pressure**<br>**(kPa)** | **IAC**<br>**(Wcm<sup>-2</sup>)** |
| --- | --- | --- | --- | --- | --- |
| 3 | 20.1 | 12.8 ± 1.0 | 42.1 ± 3.3 | 344.5 ± 065.5 | 1.0 ± 0.1 |
| 4 | 35.8 | 20.2 ± 3.2 | 49.8 ± 3.9 | 443.4 ± 084.2 | 1.4 ± 0.2 |
| 5 | 55.9 | 30.8 ± 4.2 | 49.4 ± 3.9 | 544.7 ± 103.5 | 2.2 ± 0.3 |
| 6 | 80.5 | 40.5 ± 4.0 | 46.4 ± 3.7 | 636.3 ± 120.9 | 2.9 ± 0.3 |
| 7 | 109.6 | 57.8 ± 4.3 | 45.8 ± 3.6 | 741.3 ± 140.8 | 4.2 ± 0.3 |
| 8 | 143.1 | 76.6 ± 5.6 | 47.3 ± 3.7 | 826.8 ± 157.1 | 5.4 ± 0.4 |
| 9 | 181.1 | 92.5 ± 6.6 | 44.9 ± 3.6 | 917.4 ± 174.3 | 6.5 ± 0.5 |
| 10 | 223.6 | 115.2 ± 8.1 | 45.3 ± 3.6 | 1016.2 ± 193.1 | 7.8 ± 0.6 |

Acoustic pressure generated by the focused US transducer ranged from 308 ± 58.5 kPa to 1100 ± 209.1 kPa, and the beam diameter at – 6 dB was 1.38 mm. Output power (W_OUT_) generated by the transducer ranged from 12.8 ± 1.0 mW to 115 ± 8.1 mW. The efficiency of the transducer ranged from 50.0 ± 5.0 to 64.0 ± 5.2%, with an average of 54.2%. Focal plane intensities (IAC) were measured to range between 1.0 ± 0.1 Wcm^−2^ and 7.8 ± 0.5 Wcm^−2^.

### Benchtop testing of Ultrasound-mediated Delivery Transducers in *Ex Vivo* Tissue

Previous work demonstrated that uptake of fluorescent particles by monolayers of Caco-2 cells could be enhanced by insonation^24^. Measuring the efficacy of this method in tissue is a crucial step towards its deployment *in vivo*. As in the earlier work with cells, quantum dots (QDs) were chosen as the particles used to measure uptake/delivery. Quantum dots are fluorescent semiconductor nanocrystals and are frequently used in imaging. They have broad excitation spectra, narrow emission spectra, exhibit almost no photobleaching, and have long fluorescence lifetimes^25^.

The small intestine was isolated from wild type (WT), and Apc^*Min/*+^ mice and QDs were delivered to the tissue. Apc^*Min/*+^ mice are heterozygous for mutations in the adenomatous polyposis coli gene (*Apc*) and are a well-established model of human familial adenomatous polyposis (FAP). FAP patients are also heterozygous for mutations in *Apc* and develop numerous polyps in their intestinal tract that progress to cancers if left untreated^26^. Apc^*Min/*+^ mice thus present a precancerous state. They also have reduced mucus production^27^, and any differences between WT and Apc^*Min/*+^ tissues could reflect differences in the mucus layer and/or changes associated with the epithelium in precancerous tissue^28^. The presence of QDs was compared in the samples with and without insonation and between healthy and precancerous tissues.

Successful delivery of QDs was recorded when the fluorescence emitted by the QDs was detected in the insonated area. In 11 of the 14 WT samples, fluorescence consistent with the accumulation of QDs within insonated areas was observed (Figure 2a), corresponding to a success rate of 79%. Accumulation of QDs was detectable in only 50% of the 14 Apc^*Min/*+^ samples. This observation, together with the fact that QD fluorescence was not observed after tissue samples were left in buffer (PBS) for more than 2 hours and the reduced mucus layer reported in Apc^*Min/*+^ tissue, suggested that QDs were lodged in the mucus and did not reach the epithelial cells.

**Figure 2:**
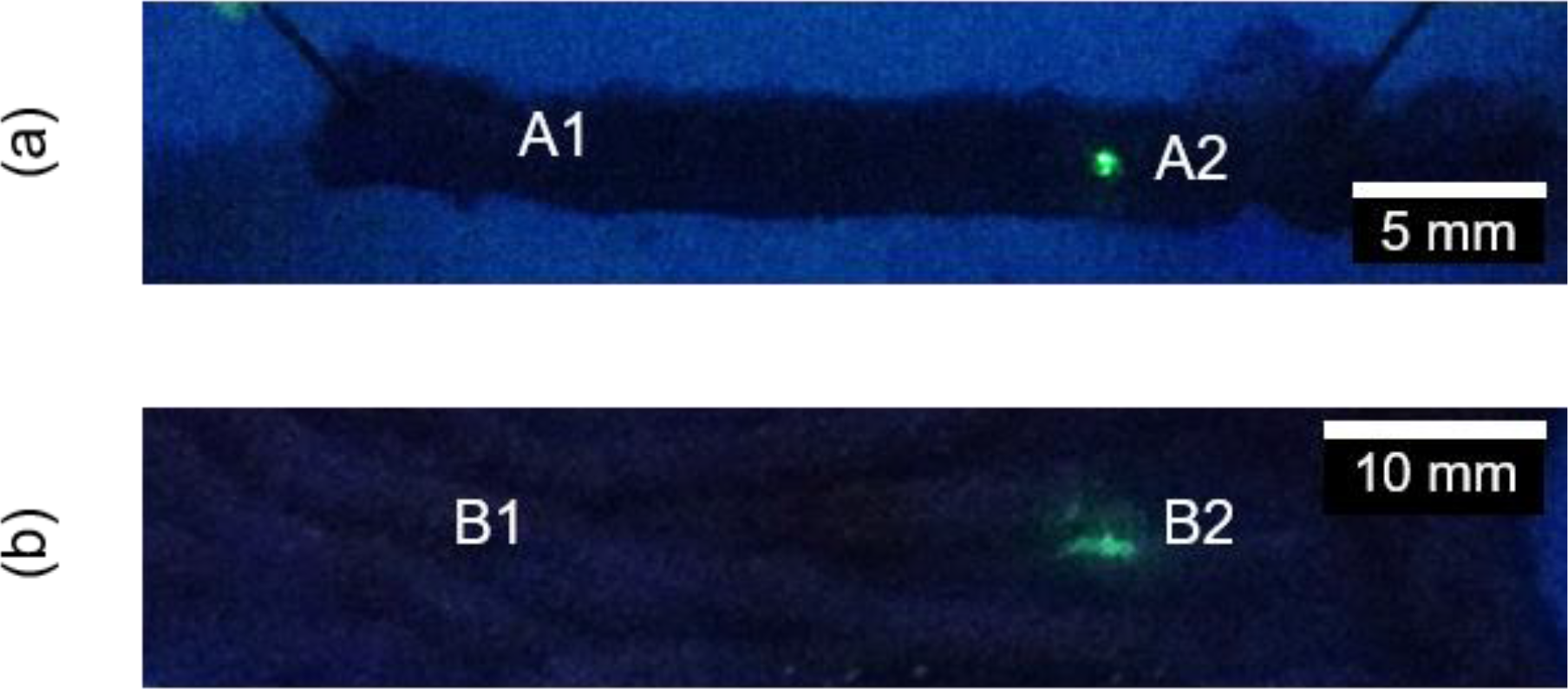
(a) Regions of wild type murine and (b) porcine intestine that were not-insonated (A1, B1) or insonated (A2, B2). In all cases tissue was exposed to Quantum dots. Quantum dots were clearly visible only when tissue was insonated (A2 and B2).

Experiments were also conducted on sections of WT porcine small intestine *post mortem*. These tissue samples were obtained and used within 20 minutes of death to minimize tissue degradation. Inspection of the porcine bowel samples exposed to QDs showed that QDs were detected only in insonated areas of bowel tissue and not in control areas (Figure 2b). Of the 16 WT porcine samples, 12 were observed to emit fluorescence within insonated areas but not control areas reflecting a success rate of 75% for *ex vivo* porcine samples.

Further murine experiments were conducted to measure the depth of penetration of the QDs in the insonated region when ultrasound was used in conjunction with MBs. Laser scanning confocal microscopy of the cross-sections of fixed murine intestinal tissue was used to determine the depth of penetration of QDs after insonation with MBs. QDs (Figure 3, green) were enveloped in the mucus layer (Figure 3, red), and but were not present in the cells in the underlying intestinal epithelial tissue (Figure 3, nuclei stained blue), demonstrating that insonation was sufficient only to drive the QDs into the mucus layer. This was further confirmed by examining a total length of sections covering 41,400 μm of insonated tissue and failing to find QDs inside cells. Processing of the tissue for staining caused the loss of much of the mucin, and only a few QDs remained with an occasional QD inside folds of villi where mucus may also have been trapped.

**Figure 3:**
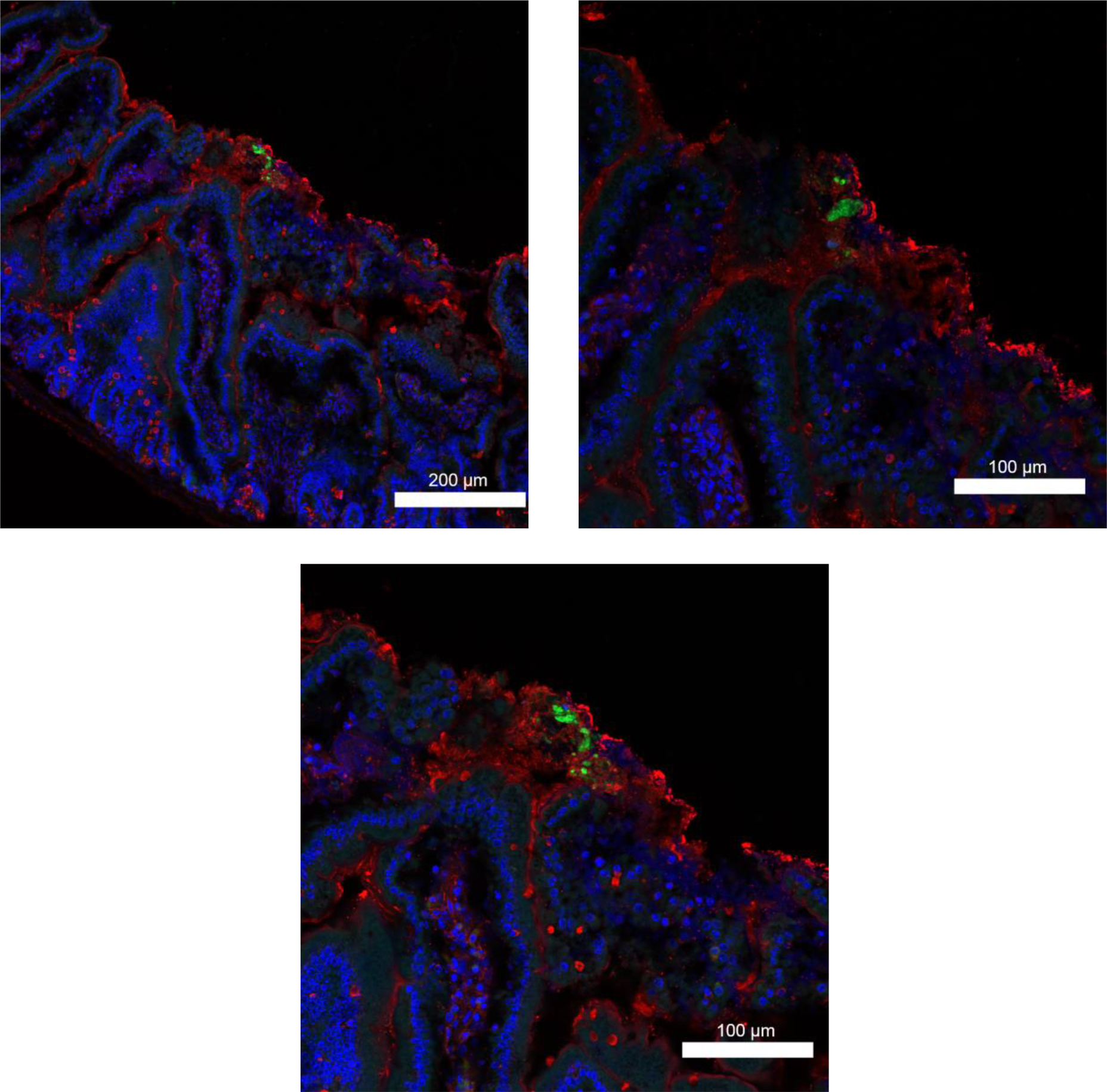
Cross-sections of different examples of murine small intestine after insonation with QDs and MBs. Images show QDs (green) are lodged in the mucosa (stained with WGA, red) after insonation and did not penetrate the underlying intestinal tissue (marked by DNA stain to show uclei, blue).

### *In vivo* Testing of Ultrasound-mediated Delivery Capsule

To determine the ability of our system to deliver reagents in live animals, five tissue samples were collected from three pigs. Two additional samples from a fourth pig were excluded because these cases, debris consumed naturally by the animal before the experimental period was found lodged in the transducer channel when the capsule was removed from the small intestine. Such debris can impede US and QD delivery and produce artificial results. As shown previously^29^, fluorescence associated with QD was detected in samples subjected to the MB/QD solution and US in the insonated regions (Figure 4a) in four of the five samples, while no fluorescence was associated with those samples subjected to just insonation (Figure 4b) or QDs (Figure 4c). Regions insonated in conjunction with MB/QD delivery were positive for fluorescence, and non-insonated controls were negative, corresponding to a success rate of 80%. High-resolution immunofluorescence imaging suggested that the QDs were lodged in the mucus layer based on the similarity between images obtained from murine and porcine benchtop trials (*ex vivo* tissue) (Figure 4d, 4e).

**Figure 4:**
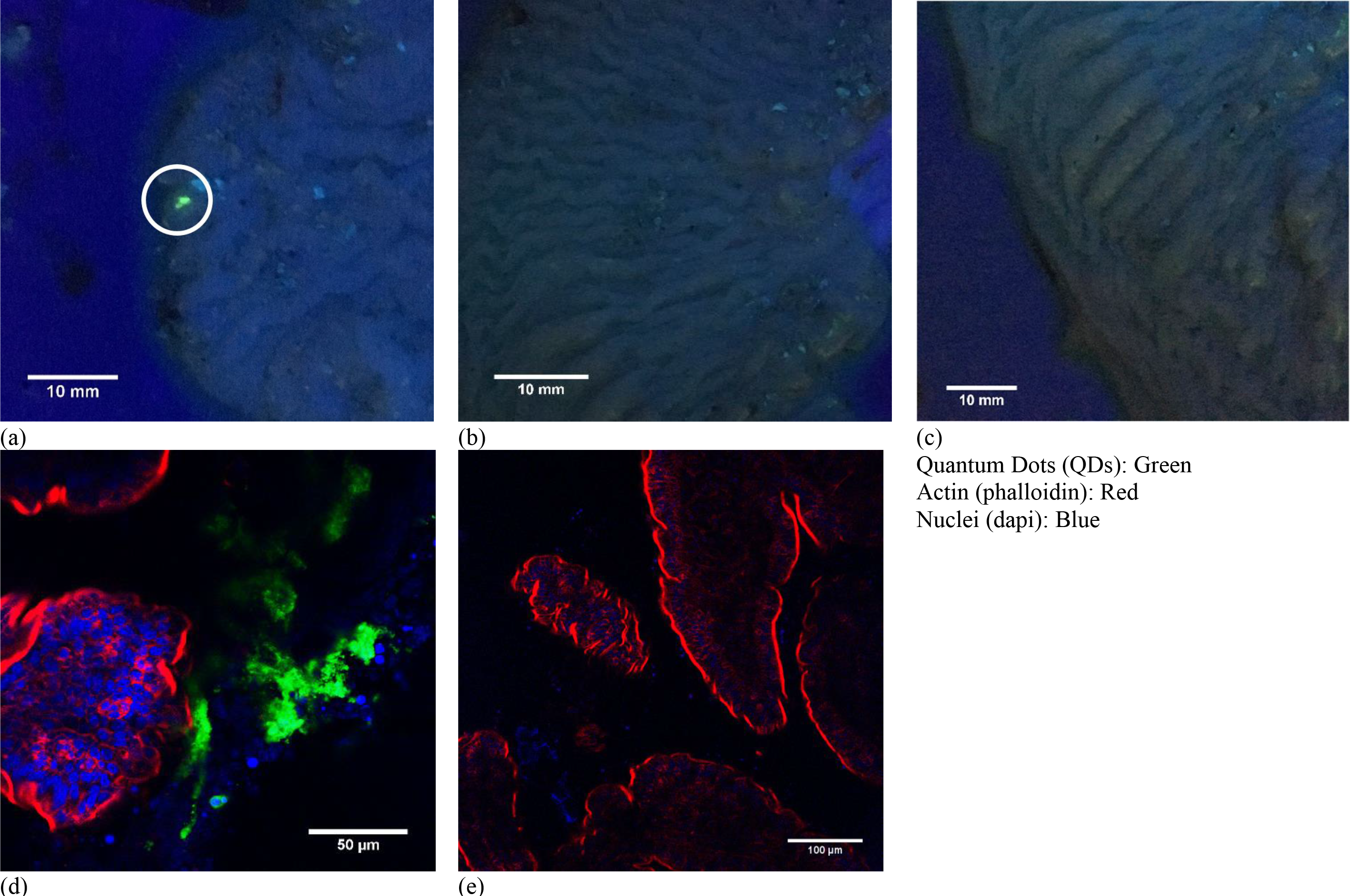
(a - c) Tissue from animal 20170921-P1. (a) Tissue sections exposed to insonation and QDs (white circle). (b) Tissue from area exposed to insonation only (no QDs). (c) Tissue from area exposed to QDs only. (d, e) Immunofluorescent staining of samples (a) and (c). The QDs were never detected inside epithelial cells marked by F-actin (red), which is highly concentrated in the apical brush border. Images a, b and c are reproduced with permission from ^29^

## Discussion

Our results provide the first demonstration of *in vivo* of ultrasound-mediated delivery of an agent in the small bowel using a proof-of-concept tethered endoscopic capsule. We attribute the higher success rate for delivery to WT murine tissue (79%) compared to Apc^*Min/*+^ tissue (50%) to the lower mucus production in the latter. Together our observations suggest that a capsule can successfully drive fluorescent QDs into the mucus of the mucosal layer of the small intestine when applying ultrasonic insonation together with MBs. The duration that the fluorescent particles persist in the mucosa will be affected by the rate of continual mucus secretion and mucosal cell shedding.

Further work is needed to determine for how long the particles remain embedded in the mucus before diffusing away and whether this residence time varies with the location along the GI tract^30^. MBs can attenuate and scatter ultrasound, making them effective as ultrasound imaging contrast agents^31^. Though it will be challenging, further work is also needed to fully characterise the relationship between drug uptake in tissue, incident ultrasonic energy and the local concentration of MBs due to their ability to attenuate and scatter the incident ultrasound. This work complements previous attempts to deliver material to the mucosa with UmTDD, which were limited to the rectum due to the size of the system employed^21^. Furthermore, this work demonstrates the potential of capsule-based, UmTDD devices for accessing the small bowel and transfer exogenous agents without requiring needles^32^, gases^33^, or other mechanisms^10^.

Delivery of therapeutics to the cells in the tissue will require UmTDD to facilitate transit through the mucus layer^34^. This could be achieved with mucus-penetrating particles (MPPs), which mimic essential surface properties of viruses that prevent muco-adhesion^30,35^. Combining drugs with MPPs could facilitate their passage through the mucus layer, and insonation could further enhance drug uptake into cells and/or through intercellular junctions. Similarly, the inclusion of mucolytic agents in the capsule is another alternative approach to facilitate the delivery of drugs into cells and tissue. Furthermore, the delivery of therapeutic agents in pathological areas such as inflamed tissue may be sufficient with this method due to the diminished mucus layer in these regions^36^.

Our results demonstrated that specific positions in the GI tract could be marked using focused US. Marking tissue to help target subsequent interventions to a specific site is a potentially useful approach to deliver treatment to a diseased site with a second follow-up capsule or surgically^37^. A capsule-based system could potentially locate diseased regions with microultrasound imaging^38^, and mark them with fluorescent particles in a US-mediated process. The fluorescently-marked (diseased) regions could then be readily identified during surgery or with a second capsule capable of fluorescence imaging^39,40^. Such a secondary capsule could also deliver therapeutic agents to the diseased site.

## Methods

### Ultrasound Transducer Design and Fabrication

A previously described prototype tethered capsule (Figure 1e) contained a focused US transducer, white light imaging camera, LED-based illumination, and a drug delivery channel^22–24^. This capsule was unsuitable for use *in vivo* as the tether was too short and inflexible, and the capsule could not be made biocompatible. Additionally, because of the diameter of the capsule relative to that of the porcine small bowel, the capsule will inevitably be in contact with the mucosa, which required it to be redesigned to allow the US focus to be at the mucosa and not below it, deeper in the tissue. This was achieved by recessing the transducer into the capsule, positioning the focus 1 mm from the capsule perimeter, instead of 4 mm as in previous designs. The additional space required for the recess meant that the new capsule (Figure 1d), could contain only a focused US transducer and drug delivery channel, and no camera or illumination. Importantly, the dimensions of the capsule (11 mm diameter, 3 mm length), remained comparable to those that are used clinically for visual diagnosis such as the PillCam® SB (Medtronic Inc., Minneapolis, MN, USA) and PillCam® Colon (Medtronic Inc., Minneapolis, MN, USA).

The transducer was produced using a curved PZ26 piezoceramic bowl (Meggitt A/S, Kvistgaard, Denmark) intended for US transmission. Each bowl had a stated transmission frequency, f = 4 MHz, an outer diameter of 5 mm, a radius of curvature of 15 mm, and an inner hole diameter of 1 mm. The hole accommodates the delivery channel used for introducing therapeutic agents. The bowl was contained in a case providing structural support that was created from VeroWhite material using the Object Connex (Stratasys Ltd., Eden Prairie, MN, USA) additive manufacturing system. The case was constrained to fit within the shell of an ingestible capsule, with shell dimensions no more than 11 mm in diameter and 30 mm in length. The case was designed in two parts to facilitate assembly and insertion of the transducer. The first part was the main body that provided support for the piezoceramic bowl and a supportive backing layer, with an outer diameter of 8 mm, and a length of 3 mm. The second part was a cap attached to the rear of the main body with an outer diameter of 8 mm, a thickness of 1 mm, and an inner hole diameter of 2 mm to allow the power cable and delivery tube for exogenous agents to pass through.

The silver electrode on the rear of the PZ26 bowl, as supplied by the manufacturer, was connected to the inner conductor of a coaxial cable (813-3426, RS Components, Corby, UK) with outer diameter of 1.17 mm, and length of 3.5 m, using conductive Ag-filled epoxy (G3349, Agar Scientific, Stansted, UK). The distal end of the coaxial cable inner connector was attached to the central pin of a cable-mount SMA connector (468-3075, RS Components, Corby, UK) using conductive Ag-filled epoxy which was subsequently cured in an oven at 80°C for 15 minutes. The backing layer was a very low acoustic impedance mixture of glass microbubbles (K1, 3M, Maplewood, MN, USA) and epoxy (EpoFix, Struers A/S, Ballerup, Denmark) at a mass ratio of 1:3 intended to provide physical support with minimal ultrasonic damping. The backing layer was applied to the rear surface of the PZ26 bowl inside the transducer case, and this was transferred to an oven to cure for 15 minutes at 70°C. Once cured, a hole of approximate diameter 1 mm was drilled through the backing layer using a 1 mm diameter drill bit to allow the delivery channel to pass through it. Polythene tubing (Smith Medical Ltd., Cumbernauld, Scotland, UK) with an outer diameter of 0.96 mm, an inner diameter of 0.58 mm, and a length of 3.5 m was used as the delivery channel. The delivery channel tubing was passed through the backing layer and PZ26 bowl until it was level with the front surface of the bowl and then fixed in place using EpoFix epoxy.

The second part of the transducer casing was attached to the rear of the main case using EpoFix epoxy. The coaxial cable and delivery channel were passed through the central hole in the case. The electrical ground connection was supplied by the outer connector of the coaxial cable attached to the front surface electrode of the PZ26 bowl using conductive Ag-filled epoxy. This connector was passed through a groove in the external surface of the case. The distal end of the outer coaxial cable was attached to the outside of the cable mount SMA connector using conductive Ag-filled epoxy and cured in an oven for 15 minutes at 80°C.

### Ultrasound field mapping

The spatial distribution of the US field produced by the focused US transducers was mapped using a commercial US field mapping (USFM) measurement system (Precision Acoustics Ltd., Dorchester, UK). The USFM system, as shown in Supplementary Figure 1, consisted of a needle hydrophone (Precision Acoustics Ltd., Dorchester, UK) moved throughout the acoustic field, including the focal region, to quantify the acoustic pressure distribution. The USFM system allows motion along *x, y*, and *z* axes with a resolution of 3.8 μm. The system was controlled, and data captured by a dedicated computer program produced with LabVIEW (National Instruments, Austin, TX, USA). A focused US transducer was placed facing downwards in the USFM water tank containing degassed water. The needle hydrophone that was used (Precision Acoustics Ltd., Dorchester, UK) had a sensitive area of diameter Ø = 0.2 mm and was positioned perpendicular to the transducers’ active element, and the motion step size was set to be 0.1mm, approximately 1/3 of the wavelength. During USFM measurement, the focused US transducers were driven at the central frequency by a 33210A signal generator (Keysight Technologies, Santa Rosa, CA, USA), and the input was a continuous sine wave, varied from 1 – 10 *V*_*pp*_, in 1 *V*_*pp*_ increments. A continuous sine wave was used to promote cavitation effects. The peak-to-peak output voltage (*V*_*pp*_) was recorded from the hydrophone and analyzed offline for pressure conversion using code produced with MATLAB (The MathWorks Inc., Natick, MA, USA). The USFM code also produced surface plots of the pressure distribution and calculated the beam diameter at −6 dB. The uncertainty in measurement at the appropriate frequency range is ±1.5 dB.

### Acoustic power measurement of transducers

Output acoustic power from the focused US transducers was measured using a commercial radiation force balance (RFB) (Precision Acoustics Ltd., Dorchester, UK) set up with a conical acoustic absorbing target. The power range of the system was 10 mW–100 W, and the frequency range was 1–10 MHz. The absorbing target was suspended in a tank of degassed water, and the transducer was mounted on a stand facing downwards toward the absorbing target. The transducer was lowered until the distance between the transducer and acoustic absorber was equal to the geometric focal distance of the transducer. The transducer was powered by a 33210A signal generator (Keysight Technologies, Santa Rosa, CA, USA). Due to the high sensitivity of the RFB system, it was encased in a draught shield to prevent airflow disturbances and minimize variation. The RFB was connected to a dedicated computer running LabVIEW software (National Instruments, Austin, TX, USA) for data acquisition and analysis. The program directly provides acoustic output power W_OUT_, considering factors such as frequency, transducer geometry, and water temperature when analyzing the data. The uncertainty in measurement at the appropriate frequency range is ± 7%. The linearity, efficiency, and intensity were calculated using W_OUT_. The linearity was obtained by comparing W_OUT_ measured by the RFB with the electrical input power, W_IN_, driving the transducer. W_IN_ was calculated using Equation 1.

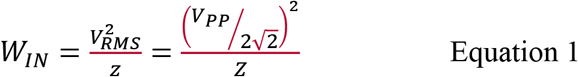

where V_RMS_ is the root mean square input voltage, and Z is the electrical impedance magnitude at the relevant frequency. The efficiency was calculated using Equation 2, where W_OUT_ is the average US output power,

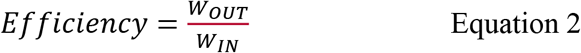

The acoustic intensity was then calculated using Equation 3:

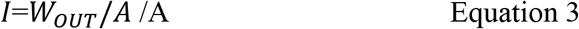

where A is the beam area at the transducer focal plane, taken at −6 dB.

### Testing of Ultrasound-mediated Delivery Transducers in *Ex Vivo* Tissue

14 CL57BL/6 wild type and 14 Apc^*Min/+*^ mice, with ages in the range of 50 - 110 days for WT (mean 85) and 55 and 95 days (mean 70) for Apc^Min/+^, with 12/14 and 10/14 female for WT and Apc^*Min/+*^, respectively, were sacrificed by cervical dislocation. All experiments involving mice were performed in accordance with UK Home Office approved guidelines and were approved by the Home Office Licensing committee (Project license P3800598E), which operates in accordance with the Animals (Scientific Procedures) Act 1986 (ASPA). The entire intestine was excised via abdominal laparotomy and immediately placed in PBS at 4°C to maintain tissue quality. The lumen was flushed with cold PBS with a syringe (Becton, Dickson and Company, Franklin Lakes, NJ, USA). The small intestine was divided into sections measuring 80–100 mm and cut along the long axis to expose the mucosa. Each sample was pinned to the acoustic absorber (shown in Supplementary Figure 2) using 25-gauge hypodermic needles (Becton, Dickson and Company, Franklin Lakes, NJ, USA), with the mucosa facing upwards. The correct orientation of the tissue was confirmed through inspection with a dissection microscope. The tissue pinned to the acoustic absorber was placed into the insonation tank (Supplementary Figure 2) and submerged in 250 ml PBS at 37°C. The tank was then transferred to the insonation system

Quantum dots (product no. 753866, Sigma-Aldrich Corporation, St. Louis, MO, USA) with a diameter of 6 nm, and an emission wavelength, *λ*_*EM*_ = 540 nm, were prepared in solution with PBS at a concentration of 100 μg/ml. The QD solution was transferred to a 5 mL syringe (Becton, Dickson and Company, Franklin Lakes, NJ, USA) and placed in the insonation system syringe driver. This QD only solution was introduced through the transducer delivery channel at a rate of 1 mL/min for 60 s. A 10 V_pp_ continuous sinusoidal waveform was applied to the transducer for 60 s per sample, producing the specific acoustic parameters presented in Table 1. Control samples consisted of a QD solution transferred onto tissue under the same conditions as above but without insonation. Additionally, the order of insonated and control samples was alternated in different experiments to account for any changes introduced by possible tissue degradation during experiments. Immediately after sonication, tissue was clamped and prepared for fixation

Additional *ex vivo* experiments were also conducted on sections of the small intestine obtained from WT pigs, where the acoustic parameters shown in Table 2 were measured to be generated by the same 10 V_pp_ continuous sinusoidal waveform. Sixteen small intestine sections were obtained from 9 pigs, aged 4 – 6 months and weighing 40 – 60 kg. Before death, the animals were used in GI experiments and killed using pentobarbital. *Post mortem* samples were obtained and used within 20 minutes to minimize tissue degradation. The small intestine section was excised via abdominal laparotomy and placed in PBS at 4°C immediately. The section was cut along the long axis to expose the mucosa and washed three times with PBS at 37°C. It was then cut into smaller sections, 50–75 mm in length and pinned to the acoustic absorber using 25-gauge hypodermic needles (Becton, Dickson and Company, Franklin Lakes, NJ, USA), mucosa facing up. The remaining length of the small intestine was stored at 4°C in PBS until insonation. The pinned tissue section was placed in the insonation tank, and a solution of 250 ml PBS at 37°C was added. The remainder of the experiment followed the protocol described for the murine samples.

Immediately post insonation/QD exposure, samples were washed with 37°C PBS using gentle agitation and rinsed using a syringe containing 37°C PBS, whilst taking care not to damage the mucosa. Samples were viewed under a 350 nm ultraviolet (UV) lamp (UVGL-58, UVP LLC, Upland, CA, USA) to assess QD uptake, and images were recorded with a digital camera. Post imaging, the method used to fix the tissue samples varied.

For immunofluorescence staining to determine the location of the QDs in murine tissue, the protocol detailed above for murine tissue was repeated but with a MB/QD solution that consisted of 5% QDs and 5% MBs in PBS. This MD/QD solution was introduced through the transducer delivery channel at a rate of 1 mL/min for 60 s. A 10 V_pp_ continuous sinusoidal waveform was applied to the transducer for 90 s per sample. Control samples were also produced whereby the MB/QD solution was provided through the transducer delivery channel at the same rate but without the presence of ultrasound. The order of control and insonated samples was varied to account for the effects of tissue degradation during the experiment. After the experiment, the small intestine samples were placed into either 4% PFA in PBS (Sigma-Aldrich Corporation, St. Louis, MO, USA) or Carnoy’s fixative, which is better able to preserve the mucus layer by fixing the mucin, and were cryoprotected overnight in a solution of 30% sucrose in PBS. The tissue was cut into 1mm pieces and placed in cryomolds before being left to incubate in 361603E OCT (VWR International, Radnor, PA, USA) for 30 minutes. The tissue was placed in the cryostat at −20°C and left to freeze before being mounted on microtome chucks using OCT. The small intestine was cut into 10-12 μm sections, with 2 to 3 sections placed onto Leica X-tra adhesive microscope slides (Leica Biosystems Nussloch GmbH, Nußloch, Germany) and left to air dry for 10 minutes and stored at −20°C. The edges of the section were blocked with a PAP pen and washed in PBS. The sections were incubated in Texas Red conjugated WGA (10 μg/ml) with Hoechst (1 μg/ml) for 60 minutes before being mounted using Vectashield anti-fade non-setting mounting agent and imaged using a Zeiss LSM 710 or LSM 880 laser scanning confocal microscopy (Carl Zeiss AG, Oberkochen, Germany). All sections were examined for QDs in the tissue. For the small intestine, two areas were sectioned and examined along the entire length of the section (area Q1 = 5,400 μm, Q2 = 22,500 μm).

### Design and Fabrication of Ultrasound-mediated Delivery Capsule

The capsule shell was designed in two parts, with pins locking the two halves together (Figure 1b). The capsule had a port to attach the tether and provide extra attachment strength. The transducer slot was at an angle such that the focus of the transducer was 1 mm radially distant from the capsule perimeter. This means that the transducer was focused on the luminal surface of the gut wall when the capsule was in contact with the wall of the GI tract^38^. The capsule was constructed in VeroWhite material using an Objet Connex 500 printer (Stratasys Ltd., Eden Prairie, MN, USA).

A tether was necessary to house the transducer power cable and the delivery channel. The tether had to be flexible enough to prevent distention of the small intestinal wall, which could potentially affect results but also be stiff enough to allow it to be used to push the capsule^23^ into the small intestine. One lesson learned from the earlier capsule is that the tether, a repurposed vascular catheter, was too stiff to be used in the GI tract *in vivo*. Instead, the tether for the present capsule was a nasoenteric feeding tube (Corpak Medsystems Inc., Alpharetta, GA, USA) with an outer diameter of 3.3 mm, an inner diameter of 2.5 mm, and length of 1.4 m. This tube was flexible but stiff enough to allow the capsule to be pushed into and along the small intestine. The tubing had a graduated scale printed on the outside and was marked every 1cm, making it possible to determine approximately how far the capsule had been inserted. This tubing allowed the capsule to be inserted up to 1.4 m into the porcine GI tract, thus leaving a further 2.0 m of an external transducer power cable and delivery channel connecting to control equipment. The extra cable ensured adequate space between the pig and measurement equipment during the *in vivo* experiments.

Medical grade epoxy (EP42HT-2MED, Master Bond Inc., Hackensack, NJ, USA) was manually dispensed by a syringe into the transducer and tether slot in one half of the capsule shell. The transducer and tether were secured in the slot (Figure 1b), and the epoxy was left to cure overnight at room temperature. The same medical-grade epoxy was also used to join both parts of the capsule shell together and left to cure overnight at room temperature. The fully assembled capsules (Figure 1c) were visually inspected for voids, and then medical grade epoxy was applied to the interfaces and left to cure at room temperature overnight.

As VeroWhite is not biocompatible, an 8 μm thick conformal coating of Parylene C was applied to the assembled capsule to not only ensure biocompatibility but also to reduce friction between the capsule and the wall of the GI tract. Parylene C was deposited using a vacuum deposition tool (SCS PDS 2010, Specialty Coating Systems, IN, USA). The surfaces were primed with A174 silane adhesion promoter before deposition. Parylene C is a USP Class VI polymer that is commonly used for coating medical devices such as surgical instruments, implants, and medical electronics^41^.

### *In vivo* Testing of Ultrasound-mediated Delivery Capsule

The performance of the capsule was measured using *in vivo* porcine models. The experiments were conducted in collaboration with the Wellcome Trust Critical Care Laboratory for Large Animals (Roslin Institute, Roslin, Scotland, UK) under license from the UK Home Office (PPL 70/8812). The experiments were approved by the Animal Welfare and Ethical Review Board of the Roslin Institute and were carried out in compliance with the terms of the Home Office license and the Animals (Scientific Procedures) Act 1986 (ASPA). Four female pigs were used with ages in the range of 3 to 6 months and weighed between 40 kg and 55 kg (Table 3).

**Table 3:** Detailed description of the four pigs used for *in vivo* delivery. Age, weight and the maximum depth of insertion achieved by the capsule through the stoma are displayed

| **Pig Identification Number** | **Age (Months)** | **Weight (kg)** | **Maximum Depth of Insertion (cm)** |
| --- | --- | --- | --- |
| 20170921-P1 | 3-4 | 40.0 | 55.0 |
| 20170921-P2 | 3-4 | 40.0 | 40.0 |
| 20171109-P1 | 6 | 55.0 | 75.0 |
| 20171109-P2 | 6 | 52.0 | 55.0 |

Anesthesia was induced with isofluorane (Zoetis Inc., Parsippany-Troy Hills, NJ, USA), vaporized in nitrous oxide and oxygen, and administered using a Bain breathing system. A cannula was inserted into the auricular vein and the trachea was intubated. Anesthesia was maintained with isofluorane. Ringer’s lactate solution (Aquapharm No. 11, Animalcare UK Ltd., North Yorkshire, UK) was administered throughout the study at 10 ml/kg/h. Normocapnia was maintained with mechanical ventilation of the porcine lungs. Vital signs were continuously monitored through the duration of the experiment by an experienced veterinary anesthetist. A stoma was created using the in-house protocol to allow direct access to the small intestine from the abdomen, bypassing the esophagus and stomach. The small intestine was flushed with PBS through the stoma. Lubrication to facilitate capsule insertion was provided by a saline drip (0.9% by weight, 1-2 drops/s) at the stoma entrance.

The transducer power cable was connected to a DG4102 signal generator (RIGOL Technologies, Beijing, China) and the capsule delivery channel was connected to a NE-1000 syringe driver (New Era Pump Systems Inc., New York City, NY, USA) containing a syringe with MB/QD solution comprising 50 μg/ml QDs (753866, Sigma Aldrich Corporation, St. Louis, MO, USA) and 1×106 MBs/mL (SonoVue MBs (Bracco Imaging, Milan, Italy) MB Ø 2 - 9 microns) in the provided physiological saline, as shown in the system diagram in Figure 5.

**Figure 5:**
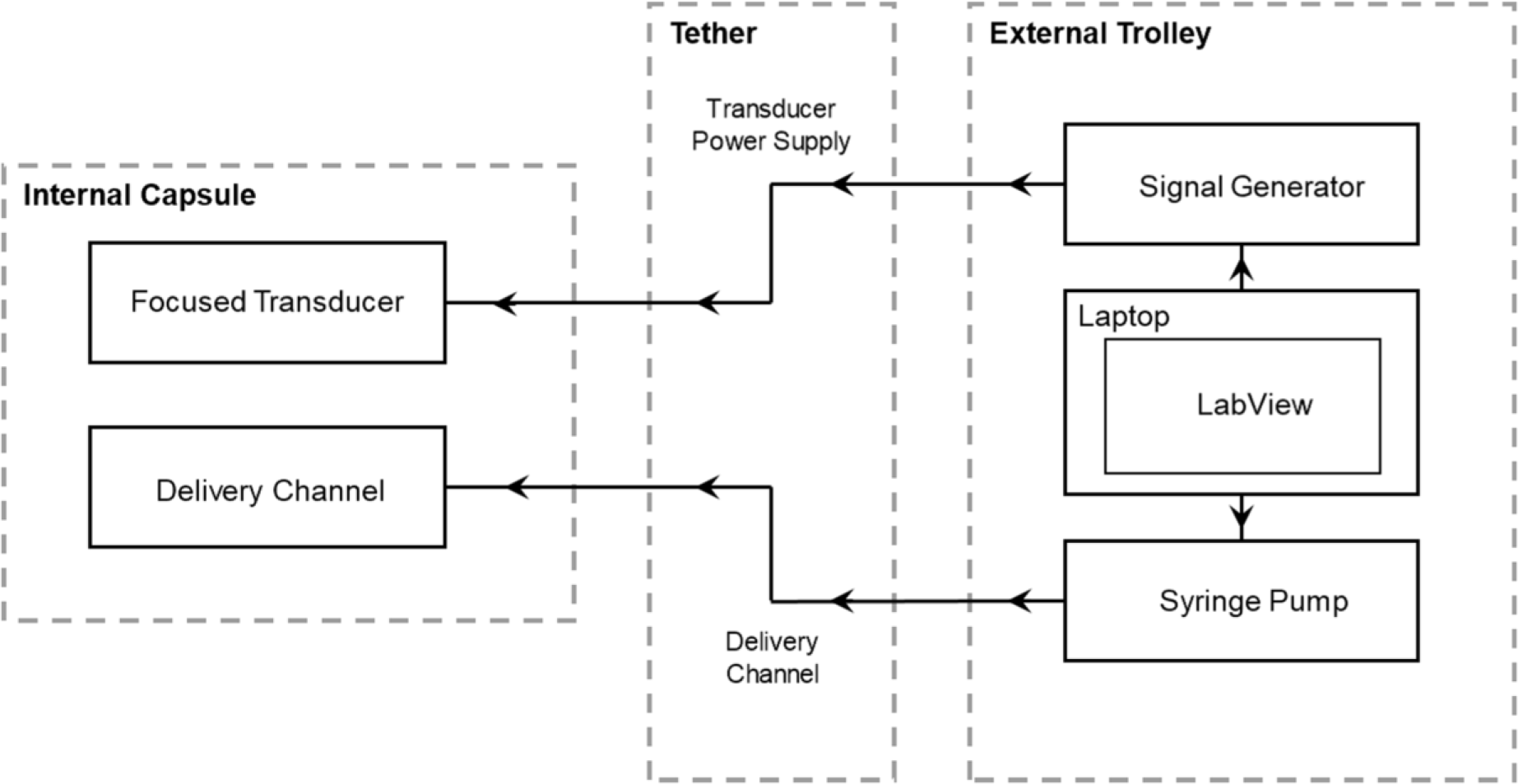
Schematic diagram of the capsule setup for the *in vivo* trial.

Each capsule was inserted through the stoma to a maximum distance at which it was difficult to manipulate the tether through the looping intestine. The distances achieved ranged from 40–75 cm (Table 3). The capsule was then removed from the stoma and cleaned with PBS. This step was necessary to assess the level of bowel preparation. A poorly cleaned bowel impeded capsule advancement and adversely affected insonation results. The capsule was primed before re-insertion by running a 2 ml MB/QD solution through the delivery channel to ensure it was not obstructed. The capsule was re-inserted into the small intestine to the maximum distance achievable and pulled back towards the stoma in positional increments of 10 cm, measured using the scale on the tether, with ‘treatment’ was delivered at each position. Different insonation/control parameters were applied at each incremental position to deliver the QDs, with the final location 10 cm from the stoma (Table 4). To test each set of parameters required 30 cm of small intestine. Therefore, the maximum number of experimental sets achievable per pig was two due to the maximum penetration depth achieved. Once all experiments were completed, the animals were euthanized while anesthetized using pentobarbital (Akorn Pharmaceuticals, Lake Forest, IL, USA).

**Table 4:** *In-vivo* experimental parameters

|  | **Ultrasound Sine Wave Parameters** |  |  | **QD Flow Parameters** |  |
| --- | --- | --- | --- | --- | --- |
|  | **Frequency**<br>**(MHz)** | **Amplitude**<br>**(V<sub>pp</sub>)** | **Duration**<br>**(s)** | **Flow Rate**<br>**(mL/hr)** | **Duration**<br>**(s, t<sub>0</sub> =**<br>**30s)** |
| Insonation | 3.98 | 8.0 | 90 .0 | 60.0 | 60.0 |
| Control | 3.98 | 8.0 | 90 .0 | NA |  |
| Control | NA | NA | NA | 60.0 | 60.0 |

### Mucosal analysis

Once death was confirmed, small intestine sections were removed via abdominal laparotomy. The sections that had been sonicated were identified by measuring the distance from the stoma. Sections were cut along the long axis and placed on a tray, mucosa facing upward. These sections were placed in PBS at 4°C immediately. The tissue was washed three times with PBS at 37°C, taking care not to damage the mucosa. The tissue was taken to a dark room and visualized under 350 nm UV light using a UV lamp (UVGL-58, UVP LLC, Upland, CA, USA). Images were acquired using a digital camera. Post-imaging, tissue samples were fixed in freshly prepared 4% PFA, pre-warmed to 37°C, for 10 minutes. After fixing, tissue was permeabilized with 1 ml permeabilization buffer (2% TX100 in PBS) in a 2 ml Eppendorf tube for 2 hours on a rocking table before washing (3 × 20 min) in PBS. 1ml of staining solution consisting of phalloidin and DAPI (Table 5) was added. PBS was added to each sample in a 2 ml Eppendorf tube wrapped in foil and left for 72 hours on a rocking table at 4°C. After staining, samples were washed 3 × 20 min in PBS on a rocking table. Samples were imaged using a Zeiss 710 confocal microscope (Carl Zeiss AG, Oberkochen, Germany) and Z-stacks were taken at the appropriate locations for each sample.

**Table 5:** List of Immunofluorescent stains used

| **Material** | **Source** | **Cat. No.** | **Excitation** | **Emission** | **Label/Dye** | **Dilution or Concentration** |
| --- | --- | --- | --- | --- | --- | --- |
| Phalloidin | Invitrogen | A12380 | 578 | 600 | Alexa Fluor 568 | 1:40 |
| DAPI | Sigma-Aldrich | D9542 | 340 | 488 | - | 1:5000 |
| WGA | Invitrogen | W21405 | 595 | 615 | Texas Red-X | 10 µg/ml |

## Acknowledgments

Financial support is gratefully acknowledged from the UK Engineering and Physical Sciences Research Council (EPSRC), Grant EP/K034537 (Sonopill Programme), and the Biotechnology and Biological Sciences Research Council (BBSRC), Grant BB/M017079/1. Microscope access was provided by the Dundee Imaging Facility; the Zeiss LSM 880 Airyscan microscope was funded by an MRC grant to the Protein Phosphorylation and Ubiquitylation Unit.

## Author contributions statement

F.S. conceived and conducted most experiments, designed and built the capsule, and analyzed results. G.C. contributed to the design, manufacturing, and assembly of the capsule as well as drafting the initial manuscript. M.T. conducted some of the experiments using mouse tissue; and contributed to the design of experiments and analysis of results. B.C. prepared the mouse and tissue for sonication, contributed to the design of experiments, conducted the *in-vivo* experiments, and analyzed the results. A.P processed, imaged, and analyzed the results of the cross-sectional imaging of murine intestinal mucosa. I.P.N. conducted preliminary murine experiments, helped with data acquisition from murine studies, and contributed to the design and running of those experiments. E.C contributed to the design of the *in-vivo* experiments and conducted them. M.Y.P.D contributed to the writing of the manuscript and provision of resources for the manufacturing and assembly of capsules. M. Thanou contributed to the supervision and planning of all aspects of this work. HM contributed to the planning and supervision of microbubble experiments and UmTDD delivery, provision of equipment, and discussion of the results. SC contributed to the supervision and planning of all aspects of this work, analysis of the experimental results, and provision of resources required for the experiments. I.N. contributed to experiments, the writing of the manuscript, provision of resources for biological experiments, and supervision of all aspects of this work. All authors reviewed the manuscript.

## Additional information

The authors declare no competing financial interests.

