## Supplementary Information for "Ultrasound Mediated Delivery of Quantum Dots from a Capsule Endoscope to the Gastrointestinal Wall"

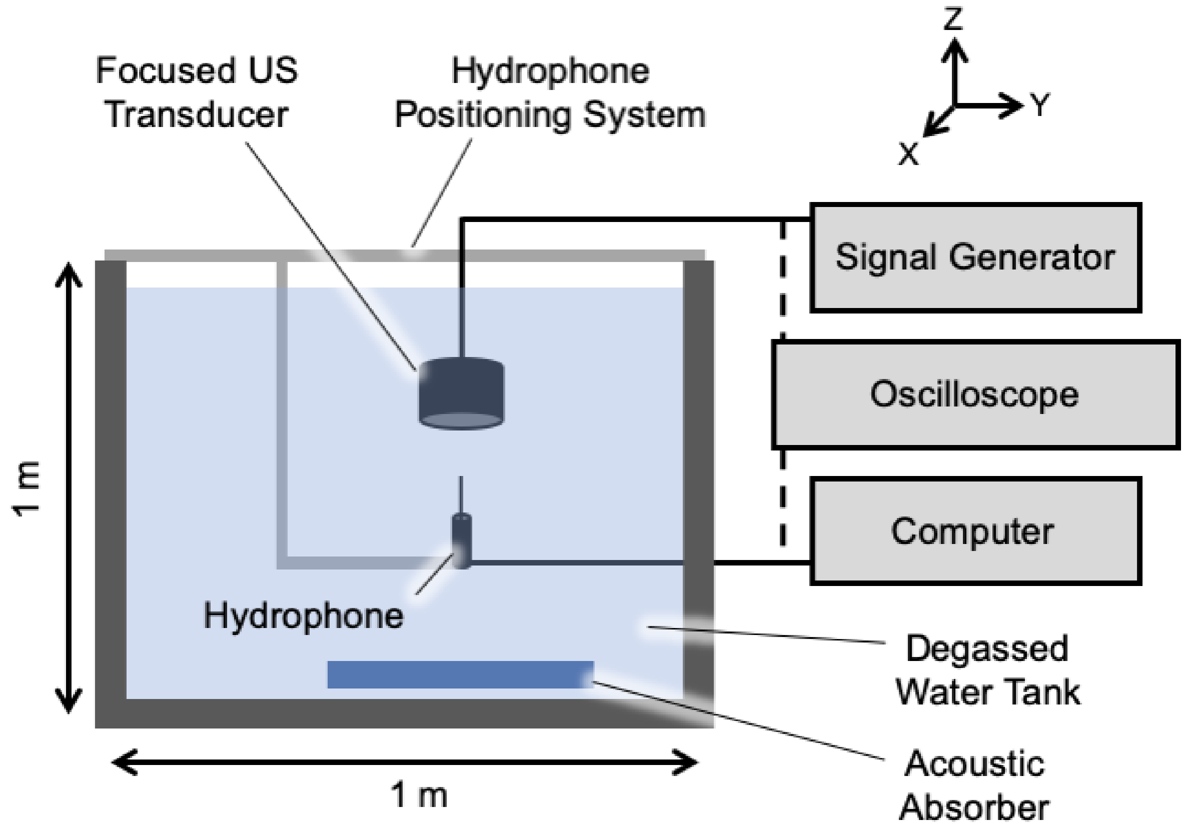


Supplementary Figure 1: Configuration of the ultrasound field mapping system with the needle hydrophone in the degassed water tank, centered and facing the active surface of the transducer.

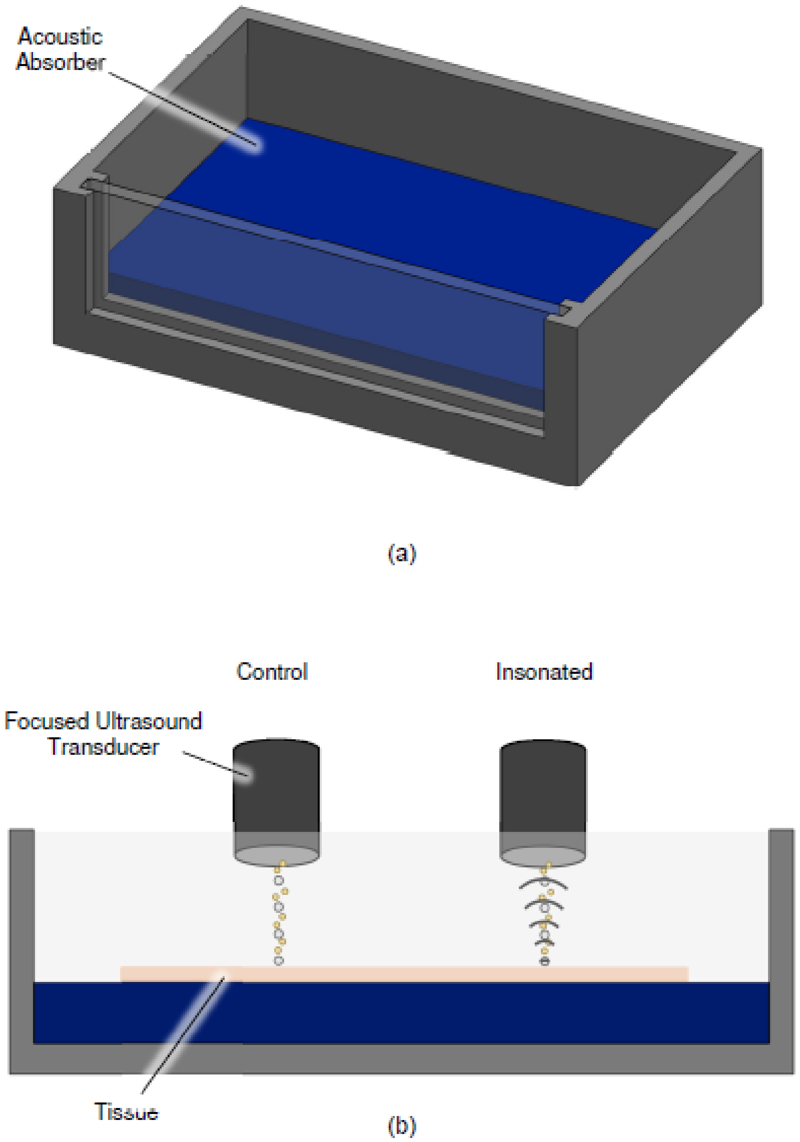


Supplementary Figure 2: (a) Insonation tank constructed with the same footprint as a standard multiwell plate. (b) Tissue was pinned to the acoustic absorber and the tank was filled with PBS. Experiments applied QDs without insonation (control, left), and QDs with insonation.

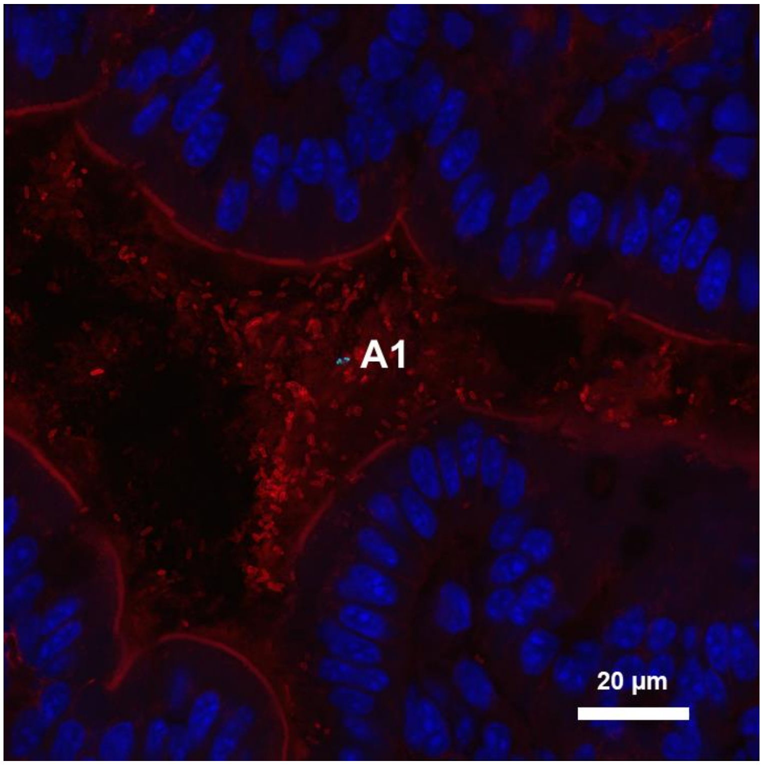


(a)

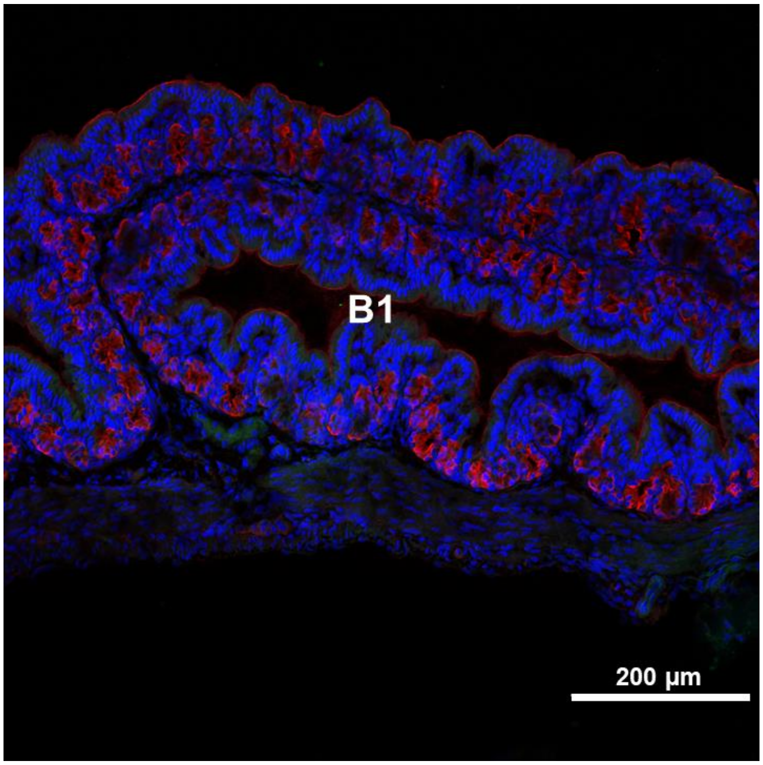


(b)

Supplementary Figure 3: Images showing staining of the murine colon and minimal presence of QDs. Images show errant QDs (green) are present on top of the mucosa layer (stained with WGA, red) at A1 and B1 and did not penetrate the mucosa or the underlying intestinal tissue (marked by DNA stain to show nuclei, blue). Scale bars represent 20 μm in (a) and 200 μm in panels (b).
